# Mass Spectrometric Identification of SARS-CoV-2 Proteins from Gargle Solution Samples of COVID-19 Patients

**DOI:** 10.1101/2020.04.18.047878

**Authors:** Christian Ihling, Dirk Tänzler, Sven Hagemann, Astrid Kehlen, Stefan Hüttelmaier, Andrea Sinz

## Abstract

Mass spectrometry (MS) can deliver valuable diagnostic data that complements genomic information and allows us to increase our current knowledge of the COVID-19 disease caused by the SARS-CoV-2 virus. We developed a simple, MS-based method to specifically detect SARS-CoV-2 proteins from gargle solution samples of COVID-19 patients. Our protocol consists of an acetone precipitation and tryptic digestion of proteins contained within the gargle solution, followed by a targeted MS analysis. Our methodology identifies unique peptides originating from SARS-CoV-2 nucleoprotein. Building on these promising initial results, faster MS protocols can now be developed as routine diagnostic tools for COVID-19 patients.

**Graphical Abstract:** Image credit (left): Gerd Altmann, Pixabay License, https://pixabay.com/illustrations/corona-coronavirus-virus-covid-19-4959447

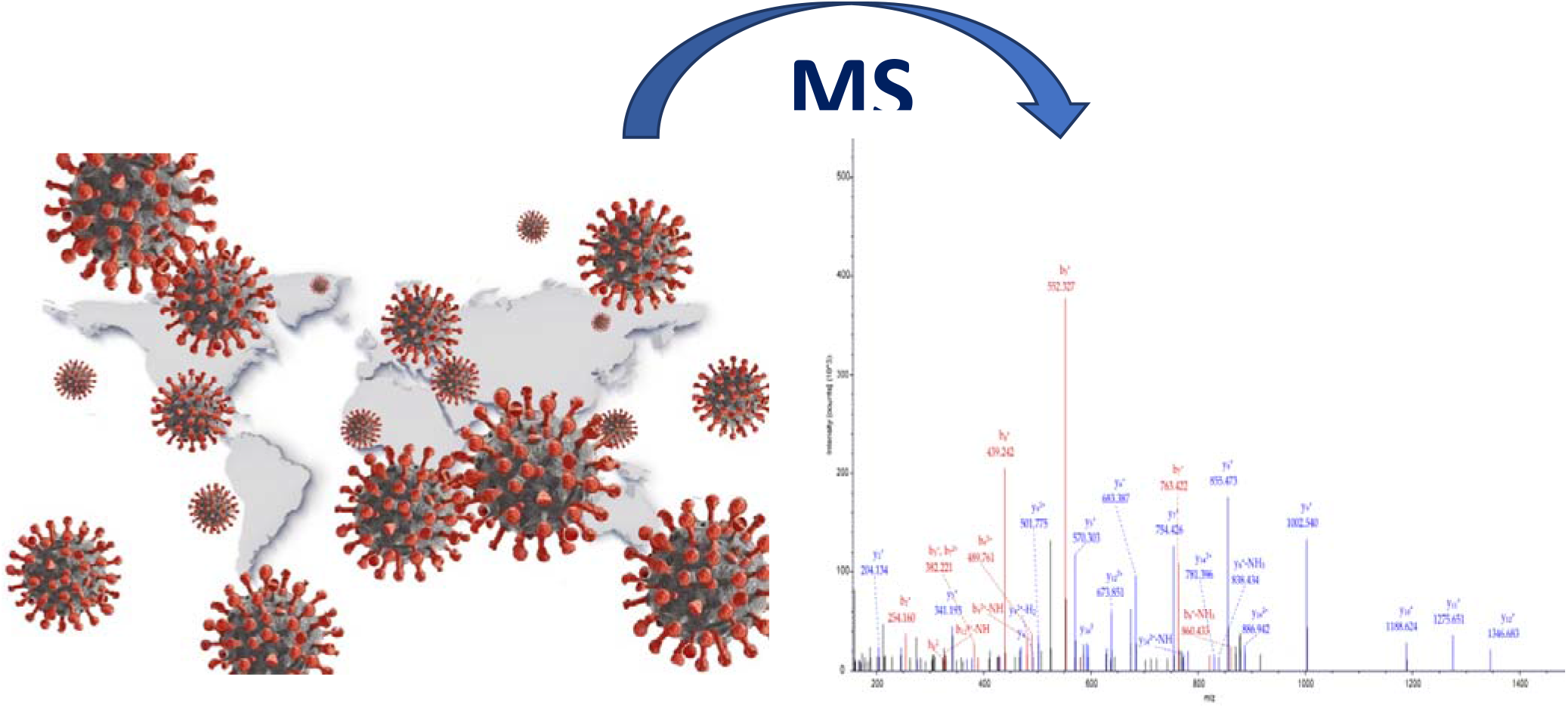

## Introduction

The disease COVID-19, caused by the SARS-CoV-2 virus, was declared pandemic by the World Health Organization on March 12, 2020 [1] and has caused more than 130,000 fatalities worldwide (date April 16, 2020). Providing novel mass spectrometry (MS)-based diagnostic tools that complement genomic approaches is one of the major goals of the recently formed COVID-19 mass spectrometry coalition (www.covid19-msc.org) [2]. Based on the prospective goals of this coalition, the aim of this “proof-of-principle” study is to highlight the potential of MS in identifying SARS-CoV-2 proteins, even from highly diluted samples, such as gargle solutions of COVID-19 patients.

## Experimental Section

### Protein Precipitation and Digestion

Gargle solutions (20 ml in 0.9% NaCl) were obtained from three patients with confirmed COVID-19 infection. All samples had been classified as SARS-CoV-2-positive by three reverse transcription and quantitative polymerase chain reaction (RT-qPCR) analyses identifying E-, S- and N-gene RNAs. For protein precipitation, 1 ml of acetone (−20°C) was added to 750 μl of gargle solution and stored overnight at −20°C. After centrifugation, the supernatant was discarded and the pellet was dissolved in 10 μl of SMART Digest buffer (Thermo Fisher Scientific). The solution was kept at 75°C for 10 min. Afterwards, 1 μl of PNGase F (non-reducing, NEB) was added to remove protein *N*-glycosylations and the sample was incubated at 50°C for 10 minutes. 1 μg of trypsin (SMART Digest, bulk resin option, Thermo Fisher Scientific) was added in 50 μl of SMART Digest buffer (Thermo Fisher Thermo Fisher Scientific) and the solution was incubated at 70°C for 30 min. Reduction and alkylation were performed with dithiothreitol (DTT) and iodoacetamide, respectively. Before LC/MS analysis, trifluoroacetic acid (TFA) was added to a final concentration of 0.5% (v/v).

### Nano-HPLC/Nano-ESI-Orbitrap-MS/MS

LC/MS analysis was performed on a nano-HPLC system (UltiMate RSLC 3000) coupled to an Orbitrap Fusion Tribrid mass spectrometer with nano-ESI source (Thermo Fisher Scientific). The samples were loaded onto a trapping column (Acclaim PepMap C18, 300 μm x 5 mm, 5 μm, 100Å, Thermo Fisher Scientific) and washed for 15 min with 0.1 % (v/v) TFA at a flow rate of 30 μl/min. Trapped peptides were eluted using a separation column (200cm μPAC, C18, PharmaFluidics) that had been equilibrated with 3 % B (A: 0.1 % (v/v) formic acid in water; B: 0.08 % (v/v) formic acid in acetonitrile). Peptides were separated with linear gradients from 3 to 10 % B over 15 min, followed by 15 % B to 30 % B over 165 minutes. The column was kept at 30°C and the flow-rate was set to 300 nl/min. Targeted MS analysis was performed based on an inclusion list of tryptic peptides from SARS-CoV-2 proteins (see Supporting Information) [3]. In case none of the target masses from the inclusion list was detected, a TOP-5s method was applied. The resolving power in MS mode was set to 120,000 at *m/z* 200. A parallel data acquisition strategy was employed, relying on Orbitrap-HCD (higher energy collision-induced dissociation, R = 15,000) and linear ion trap-CID (collision-induced dissociation)-MS/MS experiments. Database search (UniProt human, dated November 2019 with 20,315 entries, and SARS-CoV-2 [3]) was performed with Proteome Discoverer version 2.4 (Thermo Fisher Scientific). The search was restricted to human and SARS-COV-2 proteins; two missed tryptic cleavages were allowed, carbamidomethylation of Cys was set as fixed modification, while Met oxidation and deamidation of Asp, due to PNGase F treatment, were set as variable modifications.

## Results and Discussion

Protein MS analysis of the three gargle samples from COVID-19 patients revealed a vast abundance of human proteins (Supporting Information). Strikingly, our protocol allowed us to identify unique peptides originating from SARS-CoV-2 nucleoprotein that were identified in two out of three samples (Figure 1). These results are in agreement with preceding PCR analyses, exhibiting a higher viral load in the respective samples. A peptide originating from SARS-CoV-2 nucleoprotein, comprising the sequence RPQGLPNNTASWFTALTQHGK (amino acids 41-61; Figure 2), was found in both samples giving first hints to its value as a potential biomarker for an MS-based COVID-19 diagnostic method. SARS-CoV-2 nucleoprotein packages the positive-strand viral genome RNA into a helical ribonucleocapsid and plays a fundamental role during virion assembly through its interactions with the viral genome and membrane protein M [3]. It is also important for enhancing the efficiency of sub-genomic viral RNA transcription, as well as viral replication [3]. The N-gene was found to be the most expressed viral gene [4], which is in agreement with peptides of SARS-CoV-2 nucleoprotein being detected in our MS analysis. Clearly, further experiments are needed to test larger patient cohorts, but our findings show that SARS-CoV-2 proteins can be identified from highly diluted solutions. Our current method has the drawback of long analysis times (3 hours per sample), requiring significant optimization to be applied to high-through-put measurements of patient samples. Also, more sensitive mass spectrometric equipment will be required to allow the detection and quantification of a larger number of virus proteins.

**Figure 1:**
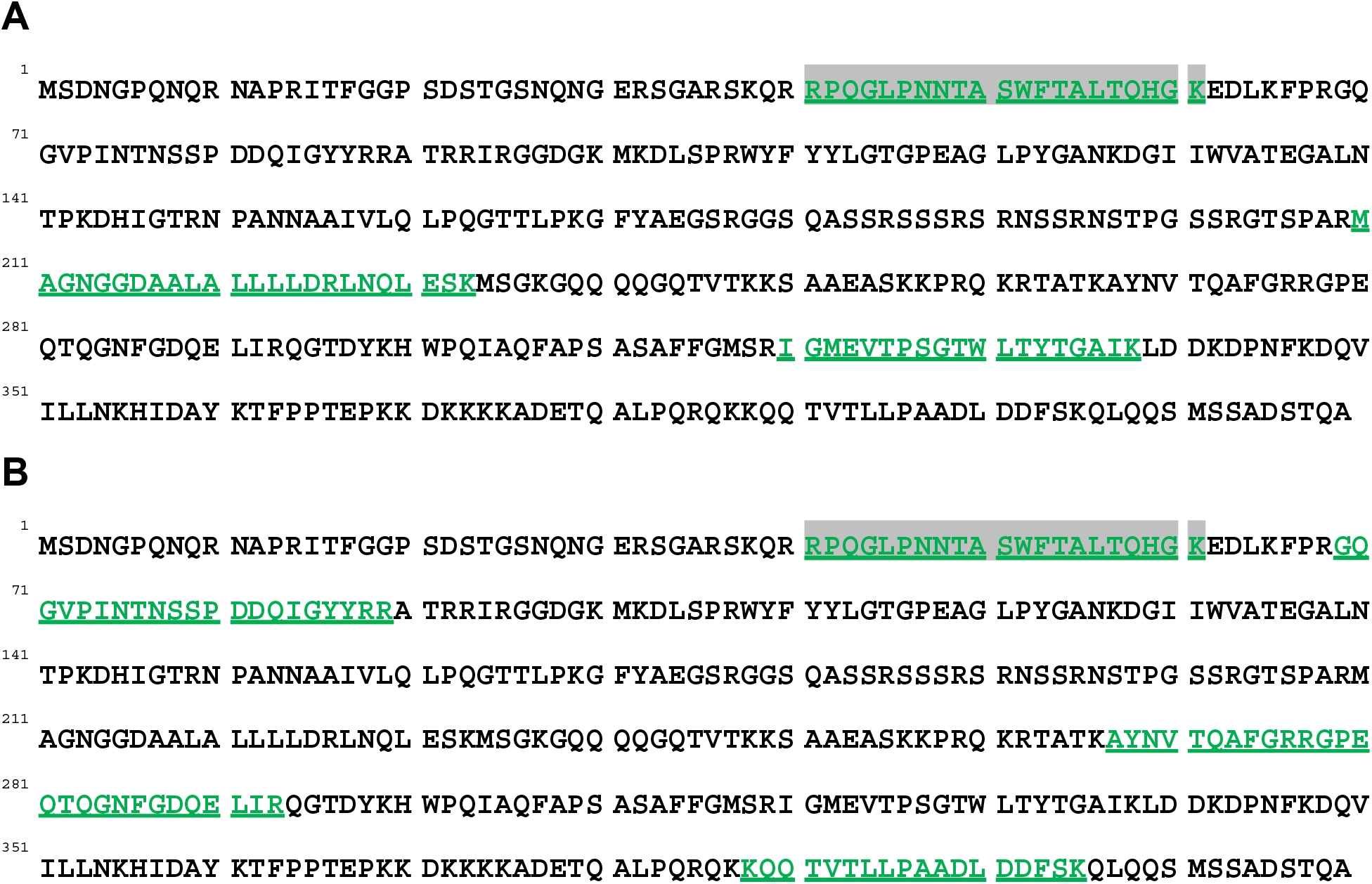
Sequence coverage of SARS-CoV-2 nucleoprotein in gargle samples (A and B) of two COVID-19 patients. Identified peptides are shown in green; the peptide (aa 41-61) identified in both samples is additionally highlighted in grey.

**Figure 2:**
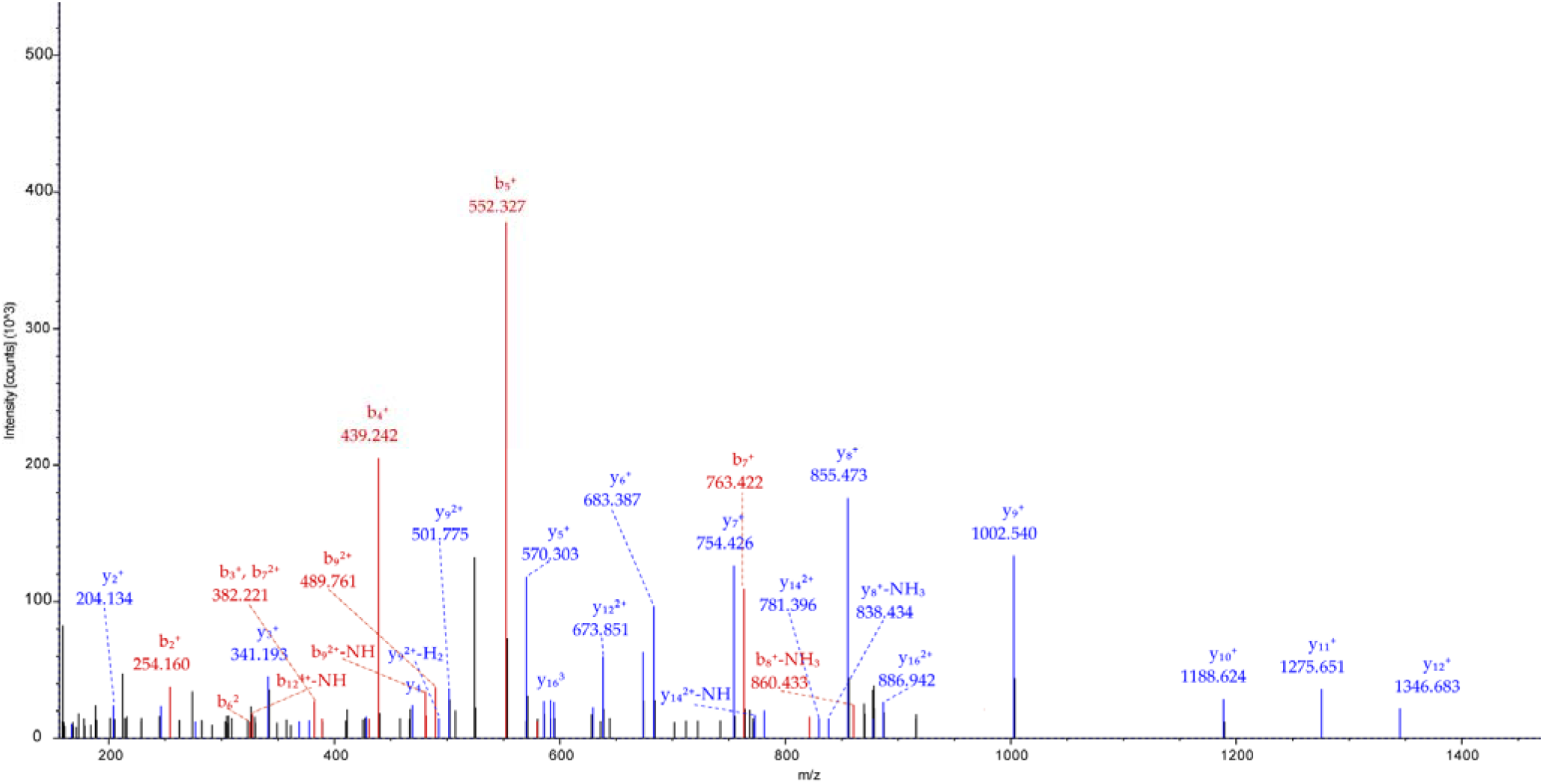
Fragment ion mass spectrum (HCD-MS/MS) of the 4+ charged signal of peptide RPQGLPNNTASWFTALTQHGK (amino acids 41-61) from SARS-CoV-2 nucleoprotein.

## Conclusions

We present a protein MS-based method to specifically detect SARS-CoV-2 virus proteins from highly diluted gargle solutions of COVID-19 patients. This straightforward method relies on acetone precipitation of proteins, followed by a tryptic digestion and a targeted-mass spectrometric analysis of SARS-CoV-2 proteins. Using this approach, we were able to identify peptides originating from SARS-CoV-2 nucleoprotein in gargle solution samples. We are planning to apply our protocol to additional sample material, such as bronchoalveolar lavage. The future goal is to provide robust, sensitive, and reliable MS-based methods as a routine diagnostic of COVID-19 patients that complements PCR-based methods.

## Supporting information

MS inclusion list

Proteins identified in sample 1

Proteins identified in sample 2

Proteins identified in sample 3

## Acknowledgements

A.S. acknowledges financial support by the Deutsche Forschungsgemeinschaft (DFG, German Research Foundation), RTG 2467, project number 391498659 “Intrinsically Disordered Proteins – Molecular Principles, Cellular Functions, and Diseases”. Prof. Gary Sawers is acknowledged for critical reading of the manuscript.

## Notes

### Competing Interest Statement

The authors have declared no competing interest.

